# Enteric Populations of *Escherichia coli* are Likely to be Resistant to Phages Due to O-Antigen Expression

**DOI:** 10.1101/2023.11.08.566299

**Authors:** Brandon A. Berryhill, Teresa Gil-Gil, Kylie B. Burke, Jake Fontaine, Catherine E. Brink, Mason G. Harvill, David A. Goldberg, Jarreth N. Navas, Kerrie L. May, Marcin Grabowicz, Konstantinos T. Konstantinidis, Bruce R. Levin, Michael H. Woodworth

## Abstract

Metagenomic data provide evidence that bacteriophage (phage) abound in the enteric microbiomes of humans. However, the contribution of these viruses in shaping the bacterial composition of the gut microbiome and how these phages are maintained remains unclear. We performed experiments with 751 combinations of 54 *Escherichia coli* and 9 phage isolates from four fecal microbiota transplantation (FMT) doses and 5 laboratory phages as samples of non-dysbiotic human enteric microbiota. We also developed a mathematical model of the population and evolutionary dynamics of bacteria and phage. Our experiments predict that as a consequence of the production of the O-antigen, most of the *E. coli* in the human enteric microbiome will be resistant to infections with the array of co-occurring phages. Our modeling suggests that phages are maintained in these enteric communities due to the high rates of transition between the resistant and sensitive states resulting from O-antigen production or spontaneous O-antigen loss, respectively. Based on our observations and predictions from this theory, we postulate that the phage found in the human gut are likely to play little role shaping the strain composition of *E. coli* of healthy individuals. Although we only investigated *E. coli*, the mechanism of resistance described here is shared among most of the Gram-negative bacteria.

**Extended Abstract:** Evidence is provided that as a consequence of O-antigen-mediated resistance, the genetically diverse array of bacteriophage in the gut microbiome of humans play little or no role in determining the densities and distribution of genetically diverse strain *E. coli* in this habitat. Our mathematical model predicts, and our experiments support the hypothesis that the phage present in the gut microbiome are maintained by replication on the minority of sensitive bacteria generated by the leakiness of O-antigen-mediated resistance.

## Introduction

The human microbiome is composed of trillions of microbes, including bacteria, viruses, fungi, protozoa, and archaea (1). The composition of the enteric microbiome has been tightly linked to human health and well-being (2–4) as well as many prevalent diseases such as cardiovascular disease, malignancy, malnutrition, and antimicrobial resistance (5–11). Interventions to modify the enteric microbiome may provide new tools to address many of these threats. However, optimal microbiome treatment strategies remain undefined. The answer to these questions requires an understanding of the factors that determine both the composition and diversity of microbes in this habitat. For the enteric microbiome, bioinformatic data has identified not only genetically diverse arrays of *E. coli,* but also genetic diversity amongst the viruses that prey on these bacteria, i.e., their bacteriophages (phages) (12). However, the contribution of these predatory viruses in determining the densities and distribution of species and strains of the bacteria in the human enteric microbiome is largely unknown.

For phages to play a role in determining the densities and diversity of *E. coli* in the gut microbiome, the dominant populations of *E. coli* in this habitat must be susceptible rather than immune or resistant to the co-existing phage. There are many mechanisms by which bacteria can become immune to phage. In phage immunity, phage adsorb and inject their genetic material into the bacteria but do not replicate; this is distinct from phage resistance. In phage resistance, bacteria prevent the phage from adsorbing in the first place. There are also several ways bacteria can become resistant to phage. The best studied of these mechanisms is envelope resistance, where because of modification of the receptor sites to which the phage adsorb, the bacteria are resistant to infections with these phage (13). Similar to this mechanism – and most relevant for this study – the O-antigen component of the lipopolysaccharides (LPS) of Gram-negative bacteria can mask the receptor sites of a number of phages, making the bacteria at least partially resistant to phages (other than those phage that use the O-antigen as a receptor) (14–17).

The results of our experiments and analysis of the properties of a mathematical model make two predictions. Firstly, as a consequence of O-antigens, the dominant populations of *E. coli* in the human fecal microbiome are resistant to the dominant population of co-existing phages in this habitat. Secondly, the observed O-antigen-mediated resistance is leaky, meaning there is a high rate of spontaneous transition from the resistant to sensitive states, thereby generating phage-sensitive minority populations of *E. coli* that can maintain the phage in this habitat.

## Results

### A method to isolate phages from fecal samples and test for the sensitivity of co-existing *E. coli* to these phages

To ascertain the susceptibility of bacteria from gut microbiomes of humans to the co-existing phages, we used Fecal Microbiota Transplantation (FMT) doses as sources of enteric *E. coli* from non-dysbiotic hosts. FMT doses are simple suspensions of human donor stool in sterile normal saline, homogenized with a benchtop stomacher device and filtered to exclude large particulate matter. We attempt to isolate phages from these samples in three ways: first, by directly assaying for the presence of phage in these samples by spotting bacteria-free filtrates of these FMT doses onto agar lawns of a broadly phage-sensitive laboratory strain of *E. coli* and searching for zones of inhibition; second, by incubating FMT doses with Lysogeny Broth (LB) to allow the phage in the FMT doses to replicate on the *E. coli* already present in these samples and then spotting the incubated suspensions onto lawns of a broadly phage-sensitive laboratory strain of *E. coli*; third, by adding both the broadly phage-sensitive laboratory strain of *E. coli* and LB to the FMTs and plating on a laboratory strain of *E. coli*. If *E. coli*-specific phage are present in the sample spotted on the lawns, either clear or turbid zones of inhibition are generated. To obtain cells from minority populations of *E.coli* from these fecal samples, we isolated the bacteria from cultures on *E. coli-*specific (minimal lactose) agar containing different antibiotics (Tetracycline; Gentamicin; Chloramphenicol; Ciprofloxacin; Ceftriaxone; Meropenem; Azithromycin; Fosfomycin; or Colistin). The methods employed to isolate phages and *E. coli* are illustrated in Figure 1, and described in more detail in the materials and methods. *E. coli* and phage isolates in these fecal samples underwent whole genome sequencing and the diversity and relationships of these bacteria and phage to each other were analyzed with comparative genomic analyses.

**Figure 1.**
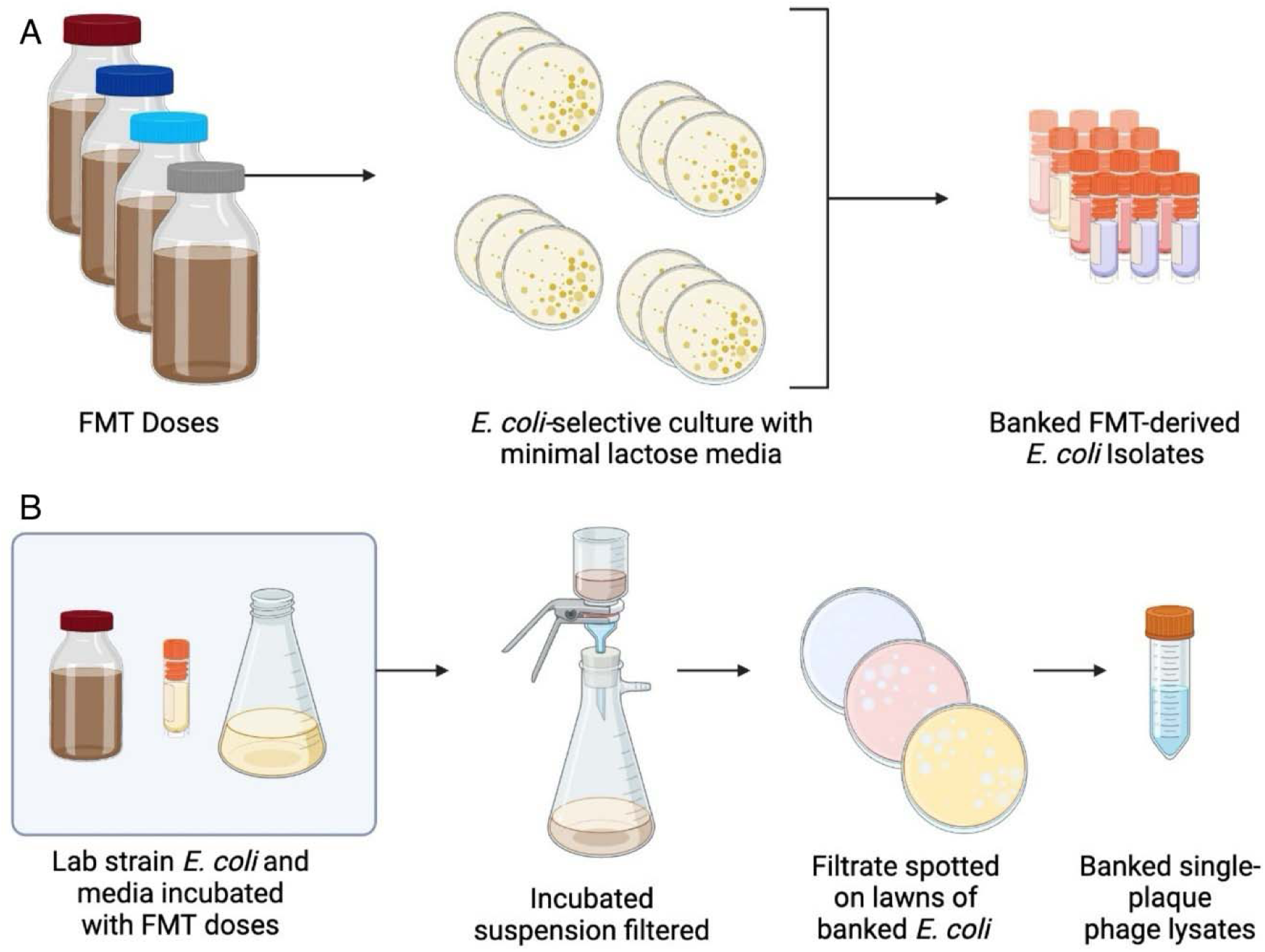
Diagram of bacteria and phage isolation protocol. (A) The method employed for isolating the dominant as well as minority populations of E. coli from the FMT samples. (B) The method employed for isolating bacteriophage from the FMT samples.

### *E. coli* and phage isolated from the FMT doses

We were not able to recover any phage by direct plating of FMT filtrates on lawns of either the laboratory strains or fecal-derived *E. coli*, indicating that the phages in these FMT samples are likely to be of low density, absent, or unable to infect and replicate on the tested *E. coli* strains. We were also not able to isolate phage after incubating the FMT doses with LB, indicating that the *E. coli* in these samples were not able to support the growth of the co-existing phages. We were able to obtain phages from three out of four of the FMTs studied by adding both the broadly phage-sensitive *E. coli* strain and LB in combination, indicating that there are low densities of phages in these fecal samples that can replicate on the lab strain of *E. coli.* The phages obtained in this way were not able to replicate on the *E. coli* isolated from these different FMTs, including almost all those from the minority of populations of *E. coli* isolated on the antibiotic plates.

Why were we able to isolate phage by adding both broth and a lab strain of *E.* coli, but not by simply adding broth? One explanation that could explain this observation is that *E. coli* in these fecal samples are already phage-resistant. We hypothesized that these FMT-derived *E. coli* are resistant to the co-existing phage and the laboratory strain is susceptibe. We reasoned this is due to expression of O-antigen in FMT-derived *E. coli* masking phage receptor sites, whereas the absence of O-antigen in the *E. coli* K-12 MG1655 laboratory strain (due to a biosynthetic mutation) makes surface receptor sites readily accessible. Two lines of evidence support this hypothesis. First, these *E. coli* isolated from FMTs bear genes coding for the O-antigen. Second, serological tests of these *E. coli* confirm the expression of O-antigens (Table 1) with one notable exception. EC1B11 does appear negative by serology yet has a predicted functional O-antigen cluster. It is known that the sensitivity of this serum assay depends on a lot of factors and it is likely that this strain just has a low expression of the O-antigen which is below the limit of detection of this assay. Neither the sequence data nor the serological tests demonstrated the existence of the O-antigen in the laboratory strains of *E. coli*.

**Table 1.** O-antigen type and production of FMT-derived E. coli isolates.

| Strain | Predicted O-antigen<br>Type | Serological Confirmation |
| --- | --- | --- |
| EC1A16 | 132 | Positive |
| EC1B11 | 1 | Negative |
| EC1B13 | 8 | Positive |
| EC1B14 | 1 | Positive |
| EC1B22 | 167 | Positive |
| EC1B24 | 1 | Positive |
| EC1C2 | 17/77/44 | Positive |
| EC1C7 | 17/77/44 | Positive |
| EC1C11 | 17/77/44 | Positive |
| EC1C12 | 17/77/44 | Positive |
| EC1C14 | 17/77/44 | Positive |
| EC1C15 | 16 | Positive |
| EC1C19 | 17/77/44 | Positive |
| EC1C20 | 17/77/44 | Positive |
| EC1C21 | 17/77/44 | Positive |
| EC1C22 | 17/77/44 | Positive |
| EC1C16 | 17/77/44 | Positive |
| EC1C18 | 17/77/44 | Positive |
| EC1D1 | 17/77/44 | Positive |
| EC1D2 | 17/77/44 | Positive |
| EC1D4 | 17/77/44 | Positive |
| EC1D6 | 74 | Positive |
| EC1D7 | 74 | Positive |
| EC1D8 | 17/77/44 | Positive |
| EC1D9 | 17/77/44 | Positive |
| EC1D12 | 74 | Positive |
| EC1D13 | 74 | Positive |
| EC1D14 | 74 | Positive |
| EC1D15 | 17/77/44 | Positive |
| EC1D17 | 17/77/44 | Positive |
| EC1D18 | 17/77/44 | Positive |
| EC1D19 | 17/77/44 | Positive |
| EC1D10 | 17/77/44 | Positive |
| EC1D11 | 74 | Positive |

### The expression of the O-antigen makes *E. coli* resistant to phage

To test the hypothesis that O-antigen production by *E. coli* mediates resistance, we performed spot assays with MG1655 (an O-antigen-negative lab strain), MG1655L5 (an isogenic mutant of MG1655 producing a medium level of the O-antigen), and MG1655L9 (an isogenic mutant of MG1655 strongly producing the O-antigen), and the phage T7. For further details on the bacterial strains and their sources see the Materials and Methods. We have restricted the assays with our constructed O-antigen strains to phage T7 for two reasons: i) this phage is known to have its receptor masked via steric inhibition due to the production and display of the O-antigen, and ii) bacteria require two genomic mutations to generate classical receptor site resistance to this phage, making the evolution of resistance unlikely during the course of our assays (18). When phage are spotted onto lawns of MG1655L9, no plaques are observed. This observation is parallel to the corresponding experiments with the *E. coli* strains isolated from FMT doses. Lawns of MG1655L5 do support the production of plaques, but those plaques are turbid rather than clear as they are on MG1655. We interpret the turbidity of the plaques found with the highly lytic phage T7 and MG1655L5 as support for the proposition of intermediate phage susceptibility.

### Accounting for the presence of phage in the apparently resistant populations of *E. coli*

If the dominant populations of *E. coli* in these fecal samples are resistant to the co-existing phage, how are the phage isolated from these samples maintained in the host enteric microbiota? Our lytic phage model (described in Supplemental Figure 1 and within the Supplemental Text section entitled “*Model of the population dynamics of resistant O-antigen bearing bacteria and lytic phage”)* illustrates that if the rate of transition from the phage-resistant (O-antigen producing) to the phage-sensitive (lacking sufficient or any O-antigen) state are sufficiently high (10^-4^ per cell per hour or higher), the phage will be maintained by growing on the minority sensitive population, the predictions of which are described in Figure 2.

**Figure 2.**
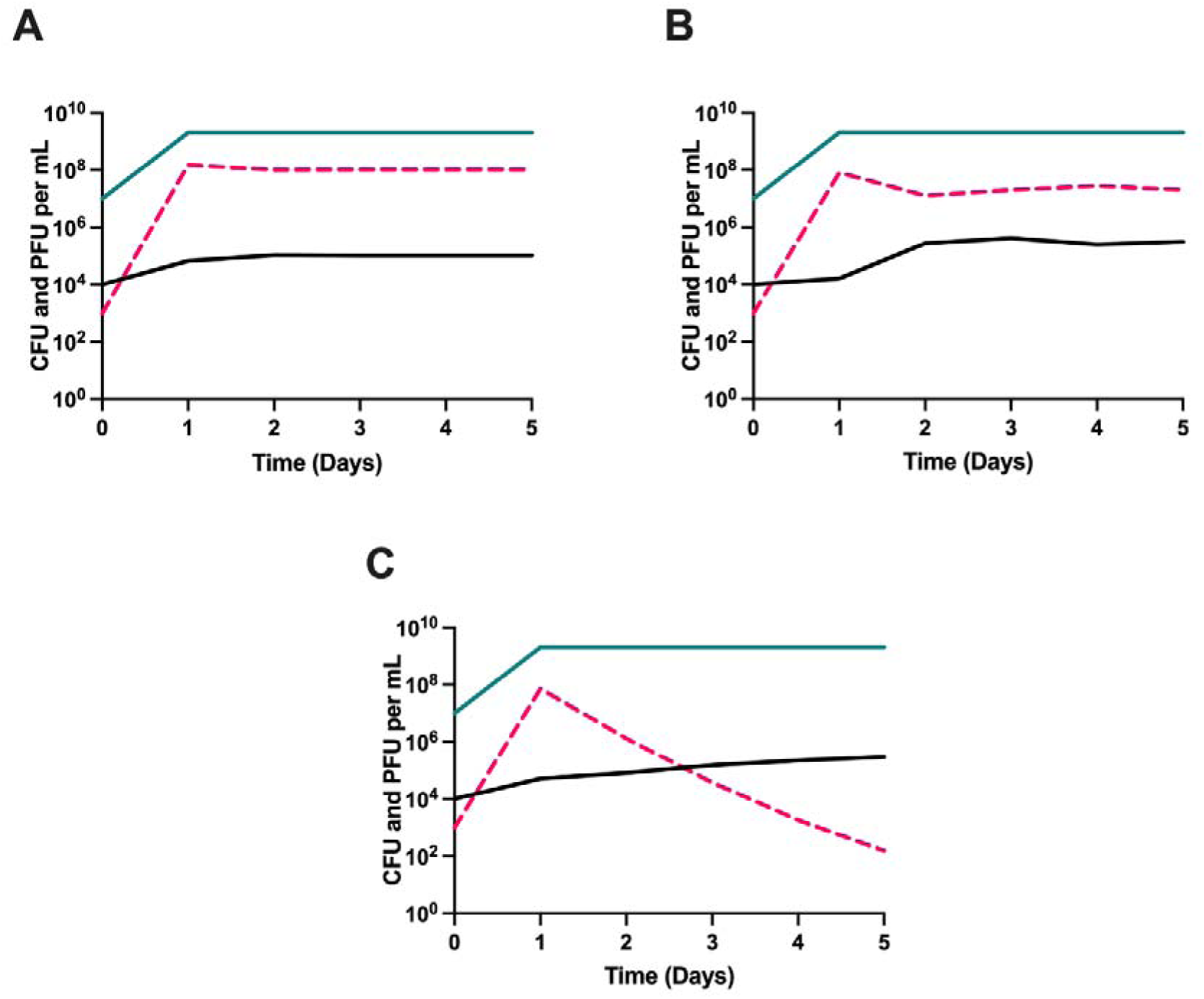
Predicted serial transfer dynamics of O-antigen-mediated resistance and a lytic phage. Simulation results showing the densities of bacteria and phage in serial transfer culture with three different transition rates between the resistant and sensitive states. At each simulated transfer, media containing 1,000 µg/mL of the limiting resource is added along with 0.01 mL of the culture at 24 hours after the previous transfer. We assume no fitness cost to resistance, and the transition rate in both directions is the same. (A) Transition rates from sensitive to resistant and from resistant to sensitive at 10-3 per cell per hour. (B) Transition rates from sensitive to resistant and from resistant to sensitive at 10-4 per cell per hour. (C) Transition rates from sensitive to resistant and from resistant to sensitive at 10-5 per cell per hour. O-antigen-expressing cells shown in green, sensitive cells shown in black, and phage shown in pink broken lines.

The model described above assumes no fitness cost for resistance, and assumes that the transition rate is the same whether bacteria are transitioning to sensitive or resistant. In Supplemental Figure 2, we consider the effects of varying fitness costs and unequal transition rates between sensitive and resistant, and find their impact to be negligible. We further explore in Supplemental Figure 3 the effects of using a temperate phage, with the model described in the Supplemental Text section entitled “*Model of the population dynamics of resistant O-antigen bearing bacteria and temperate phage”*. While the dynamics do differ slightly, the maintenance of free phage can still be explained by leaky resistance.

To empirically test this leaky resistance hypothesis, we performed serial transfer experiments with the phage T7, MG1655, MG1655L5, and MG1655L9. The results of these experiments are presented in Figure 3. In serial transfer culture, all conditions (O-antigen negative, low levels of O-antigen, and high levels of O-antigen) were able to maintain the phage. However, phage densities were significantly lower in O-antigen-expressing strains compared to wild-type MG1655. A mixed effects linear regression model showed a significant interaction of strain and serial transfer day (p=0.003) and pairwise differences in mean plaque forming unit (PFU) density for wild type MG1655 vs L5 (0.008) and wild type vs L9 (p=0.011), consistent with O-antigen-mediated resistance reducing but not eliminating phage replication. The predictions of our model, along with these experimental results, explain how phage can be maintained in populations of bacteria resistant to these viruses. This accounts for how we were only able to recover phages after adding an O-antigen-negative strain of *E. coli* and LB. It is known that loss of the O-antigen corresponds with an increase in vancomycin resistance in *E. coli* (19). Therefore, to estimate the frequency of spontaneous O-antigen loss due to leaky resistance, we isolated spontaneous vancomycin-resistant subpopulations and estimated their frequency in a culture of otherwise isogenic O-antigen expressing *E. coli* K-12. Since spontaneous vancomycin resistance can also occur through null mutations in genes of processes unrelated to O-antigen synthesis, we also used Ffm phage sensitivity to specifically diagnose loss of the O-antigen among vanomcyicn-resistant colonies. The Ffm phage receptor is the LPS core oligosaccharide which is ordinarily masked by attached O-antigen, causing resistance. Hence, the rate of both Ffm-sensitivity and vancomycin-resistance in a population can reliably diagnose lack of cell surface O-antigen. Using these measurements, we estimate that the rate of spontaneous O-antigen loss is 7.08×10^-6^ ± 6.66×10^-6^ (Fig S4). This is a low-end estimate since we know that it cannot detect mutations that truncate the LPS core oligosaccharides – these mutation would cause loss of cell surface O-antigen and vancomycin-resistance, but by destroying the Ffm receptor, would score as Ffm-resistant. Futhermore, to ensure the serial transfers results are not specific to the O-antigen serotype presented in Figure 3, we conducted parallel serial transfer experiments with one of each of the serotypes in Table 1. Parallel results obtained and are shown in Supplemental Figure 5.

**Figure 3.**
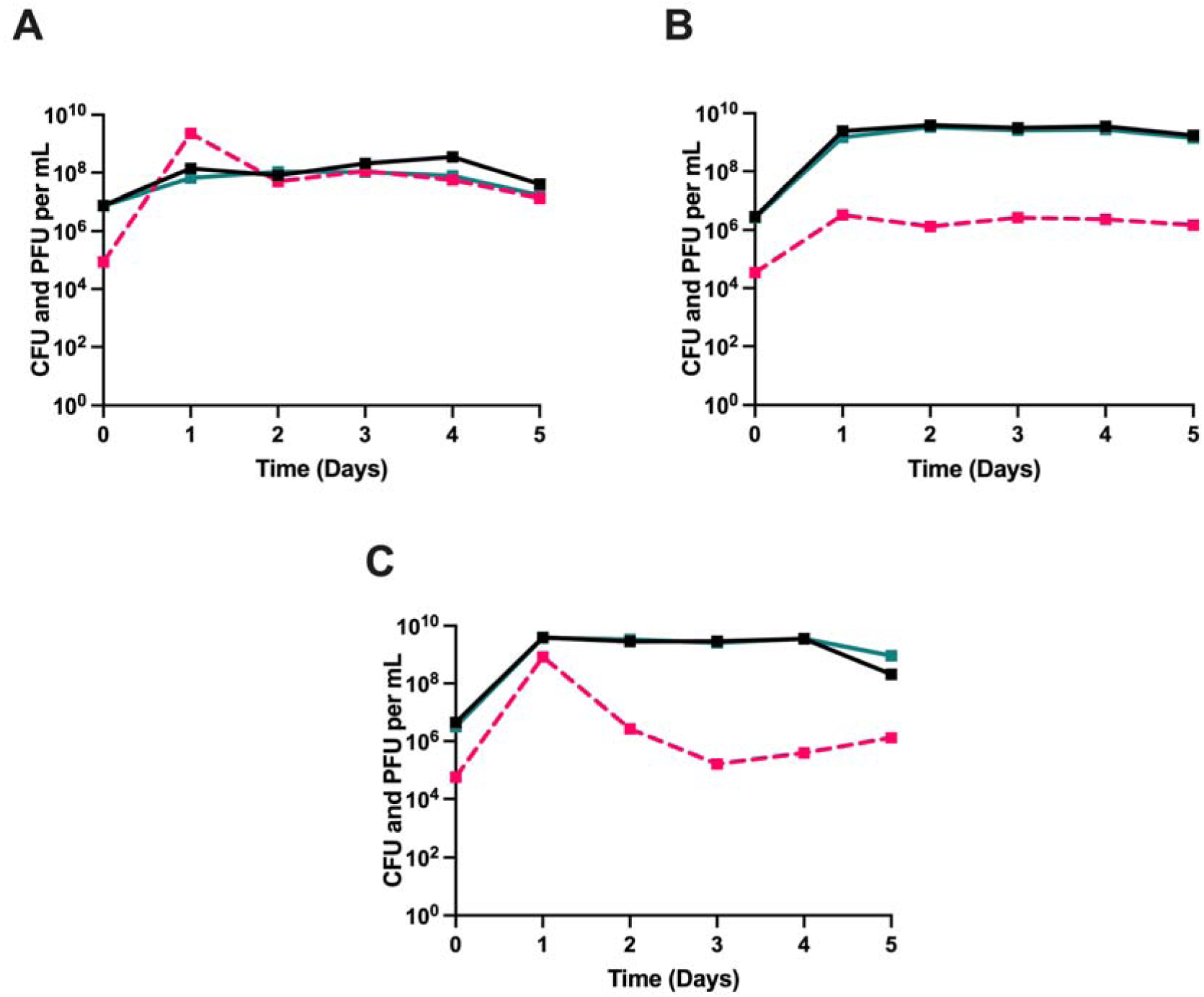
Serial transfer with phage T7 and strains of E. coli MG1655 lacking and expressing different levels of the O-antigen. Three variants of E. coli MG1655 serially transferred for five days in LB broth with the phage T7. Shown in solid lines are bacterial densities with the black line representing a phage-free control. The broken pink lines represent phage densities. Shown is one of three biological replicates. (A) Wild-type MG1655 that does not produce the O-antigen. (B) MG1655L5, which produces a medium amount of the O-antigen. (C) MG1655L9, which produces a high amount of the O-antigen. Two other, qualitatively similar, replicates are presented in Supplemental Figure 6. These results show lower but consistently measurable phage densities in E. coli populations expressing O-antigen.

### Genomic diversity of *E. coli* and its phages in FMT doses

We identified an apparent dominant and several minority *E. coli* strains in each FMT dose. These results demonstrate successful isolation of distinct populations of *E. coli* in the fecal microbiomes of several healthy donors. For each FMT donor, we assessed the diversity of recovered *E. coli* strains via alignment of predicted gene clusters, average nucleotide identity (ANI), *in silico* Multi-Locus Sequence Analysis (MLSA), and *in silico* H and O serotype prediction; we display these results in Figure 4A. We chose to predict the H and O serotype *in silico* as the diverse expression of these antigens were traditionally a measure of *E. coli* diversity (20). When applying an ANI-based genomovar definition of ≥99.5% ANI, we identified a median of 2.5 (range 2-3) culturable *E. coli* strains per donor (21). In general, classification by MLSA was consistent with the ANI-based clustering. However, strains with similar ANI and of the same MLSA displayed a greater diversity in predicted O-antigen type. These findings mirror previous studies (22) which found that healthy individuals carry diverse *E. coli* strains, confirming that our methods are readily capable of capturing the diversity in the fecal microbiome.

**Figure 4.**
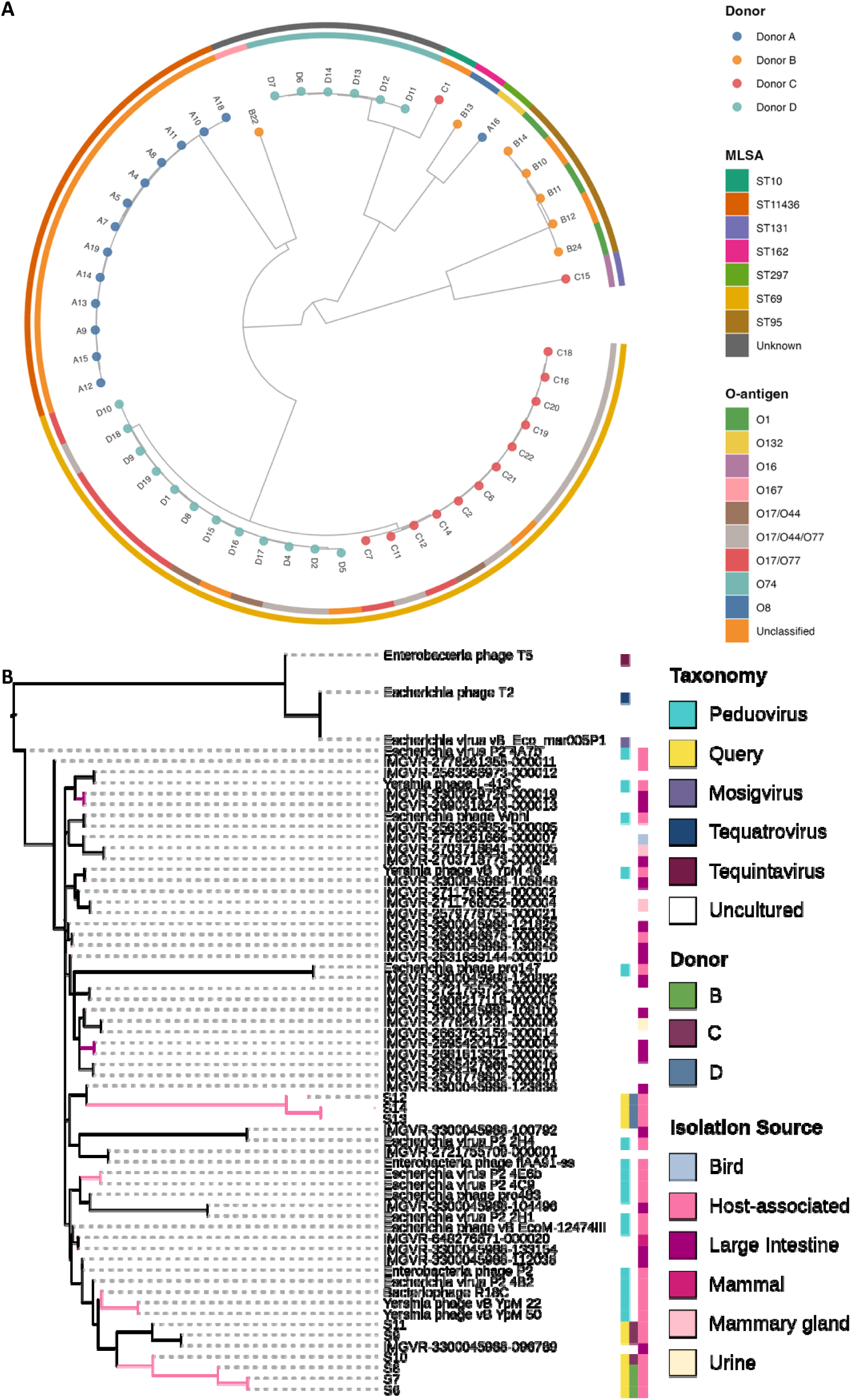
Diversity of *E. coli* and phages isolated from FMT doses. **(A)** Neighbor-joining tree constructed from pairwise average nucleotide identity (ANI) of 53 *E. coli* genomes cultured from four healthy stool donors. Tip nodes are colored by donor, inner and outer rings indicate *in silico* predicted O-antigen serotype and multilocus sequence analysis (MLSA) sequence type, respectively. Each donor harbored multiple distinct lineages differing in ANI, MLSA type, and predicted O-antigen. **(B)** Bacteriophage consensus maximum likelihood phylogenetic tree based on core genes of the phage genomes in this study and close relatives from public databases. Thirty-two genes were shared in more than 55/60 of the genomes in the major AAI cluster of Supplemental Figure 6 and were used to build individual trees as described in the Methods Section.

Of the four FMT donors, phages were recovered from three donors (Donors B, C, and D). Three unique phages were recovered from each of these three donors; thus, we recovered a total of nine distinct phages. Of note, we were not able to recover any phages from Donor A with any of the above-described isolation methods. A phylogenetic analysis of the phage genomes was conducted by generating an average amino acid identity (AAI) heatmap (Supplemental Figure 7) and consensus maximum likelihood phylogenetic trees based on the core phage genomes (Figure 4B). The reference *Peduovirus* genomes, genomes isolated in this study, and the IMG/VR genomes shared AAI values of 90%, which reflects the high similarity in gene content between these phage. The consensus tree of these gene trees, estimated using Astral (23), is shown in Figure 4B. The phylogeny demonstrates that the genomes of Donor D are only distantly related to those of B and C. The phages from Donors B and C also appear to cluster separately, with the exception of a single phage, which is closely related to the phage genomes of Donor B despite being isolated from Donor C. This suggests that each FMT donor has multiple distinct culturable phage populations and that the populations within each donor are largely distinct from the other donors.

### Susceptibility of *E. coli* isolates to phage

Our results indicate that while a high level of expression of the O-antigen is sufficient to allow phage resistance, phage are still able to be maintained by that resistant population. Furthermore, the diverse *E. coli* isolated from the FMTs were found to display a variety of O-antigen types. These observations, taken together, suggest that *E. coli* isolated from healthy human enteric microbiota are likely to be resistant to a diversity of *E. coli* phages, including those from the same community.

To get a better idea of the phage susceptibility/resistance of the *E. coli* in the FMTs we spot tested three classes of phage on lawns of each of the genetically distinct bacteria isolated from the FMTs: i) five well-characterized laboratory phages not isolated from feces; ii) the co-existing phages isolated from the same donor; and iii) phages isolated from the other donors. In total, we tested 751 combinations of 54 bacterial isolates and 14 phage isolates.

Lawns of the laboratory strain *E. coli* C lab displayed clear plaques in spot tests with the 14 tested lab and FMT-derived phages tested (Figure 5). This is not the case for lawns with the FMT-derived *E. coli* isolates, which give either no plaques (217/265, 81.9%) or turbid plaques (42/265, 15.8%). There is one exception: The phage T3 spotted on EC1D8 generated a clear plaque (0.3%). On lawns of the FMT-derived *E. coli* isolates, 392/477 (82.2%) produced no plaques and 85/477 (17.8%) produced turbid plaques.

**Figure 5.**
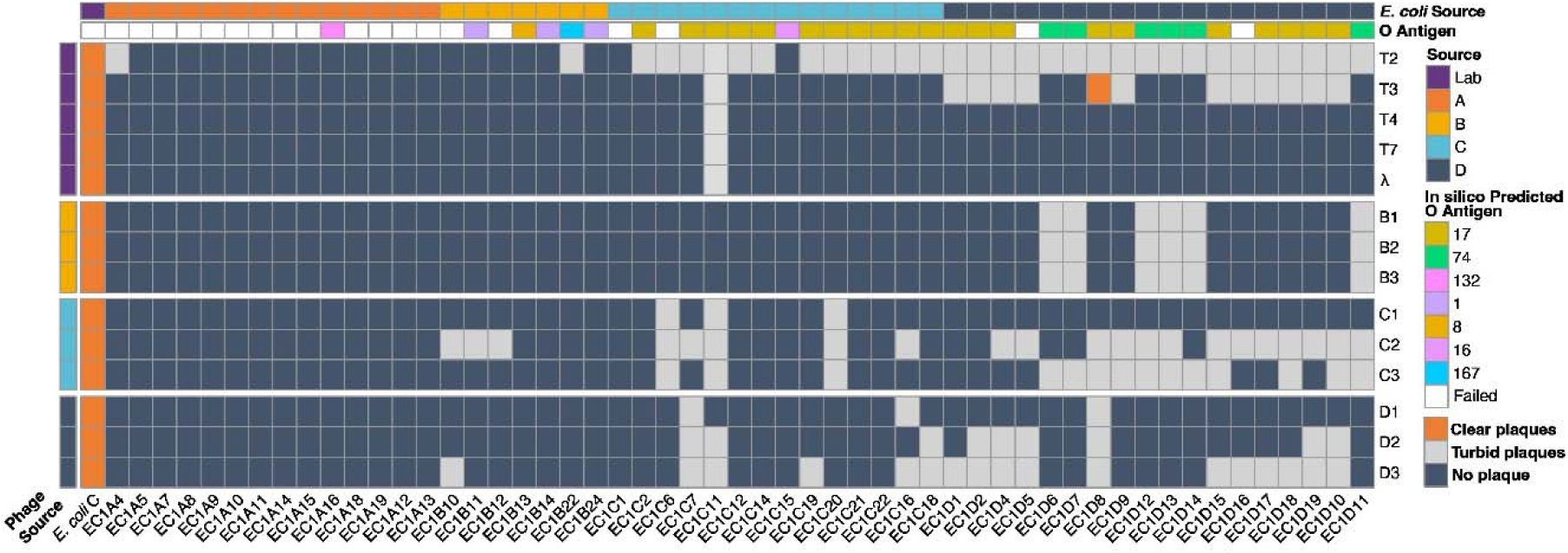
FMT-isolated E. coli do not give clear plaques when spotted with laboratory or FMT-derived bacteriophages. Well-characterized laboratory and FMT-derived E. coli strains (indicated by Source as Lab or Donor A through D) were spotted with well-characterized laboratory phages (T2, T3, T4, T7, and a virulent mutant of phage λ) and FMT-derived bacteriophages (isolated from FMT Donors B through D). Dark blue denotes no plaque formation, light grey denotes turbid plaque formation, and orange denotes clear plaque formation. Clear plaques were almost exclusively produced by the tested phages on lawns of a laboratory strain of E. coli C indicating broad resistance or immunity of FMT-derived E. coli to phages from all sources.

For phages in the FMT doses to shape the composition of a recipient’s enteric microbiota, the recipient’s *E. coli* must be sensitive to FMT-derived phage. However, when comparing combinations of FMT-derived phage and *E. coli* isolated from the same or different donors, we do not find sensitivity. We found a significantly higher proportion of turbid plaques with autologous (bacteria and phage from the same donor) phage-*E. coli* combinations compared to allogeneic (bacteria and phage from different donors) phage-*E. coli* combinations (29/117, 24.5% vs 56/360, 15.6%; chi-square p-value 0.033). Further analysis of these and the above pairs of bacteria and phage are described in Supplemental Table 1. These results, taken together, indicate that FMT-derived *E. coli* are less sensitive to phages than laboratory *E. coli* strains—not just FMT-derived phages, but phages in general. To determine if the O-antigen is sufficient to explain this generalized pattern of resistance, we obtain O-antigen-negative mutants of each serotype (Table 1) and assess their phage-suscetiblity (Supplemental Table 2). Importantly, loss of the O-antigen broadly restored phage susceptibility.

Interestingly, the expression of the O-antigen in the population is not merely coincidental. We find that the O-antigen is selected for in the presence of phage, as shown in Supplemental Figure 8. In the presence of phage T7, O-antigen expressing cells (MG1655L9) increased from ∼0.008% to 98.1% of the population within 24 hours, compared to 0.09% without phage (t-test p = 2.9 × 10⁻L). Could phage be one of the many selective pressures responsible for the ubiquity of the O-antigen in natural populations of *Enterobacteriaceae*?

## Discussion

We set out to elucidate the role of bacteriophages in shaping the composition of *E. coli* in the enteric microbiome of healthy humans. Our interpretation of these results is that bacteriophages likely play little role in determining the strain composition of co-occuring *E. coli* due to the high prevalence of phage resistance in the enteric microbiome of healthy individuals. In the case of *E. coli*, the expression of the O-antigen is sufficient to confer phage resistance while allowing for the maintenance of phage due to the random transition between a phage-resistant and -sensitive state, which we call leaky resistance. Several diverse mechanisms can convey bacteriophage resistance (such as receptor site mutation and the carriage of phage resistance genes on prophage (24, 25)) but we demonstrate that expression of the O-antigen is sufficient to account for the resistance of *E. coli* to a broad swath of phages, including those to which they have never been exposed. Our models and experiments support the hypothesis that despite this resistance, phage will be maintained in communities of these O-antigen-expressing bacteria. Moreover, we expect that this same resistance could be observed in many natural populations of *Enterobacteriaceae*, as this family broadly expresses the O-antigen. Stated another way, we postulate that resistance to co-occurring phages will be the dominant characteristic of bacteria in stable natural communities (26). This prediction is consistent with experimental studies of lytic and temperate bacteriophages *in vitro* (18, 27).

Resistant mutants almost always emerge and ascend to become the dominant population of bacteria in *in vitro* experiments with laboratory strains of bacteria and lytic phages. Nevertheless, phages are commonly maintained in these cultured populations that are dominated by resistant bacteria (27, 28). There are a variety of mechanisms by which the phage can be maintained, but their maintenance appears to play little role in regulating the densities of these bacterial cultures, and these bacterial populations are more likely shaped by other factors such as resource availability. We postulate this could also be the case in the gut microbiome. While there are many other mechanisms by which phages can be maintained, such as physical refugia as a consequence of the structure of the gut or spontaneous temperate phage induction (Supplemental Figure 3), we have demonstrated that leaky resistance is sufficient to independently account for the results observed here. There is, however, a caveat to our purported mechanism for phage maintenance in that one prediction of our model is that the density of bacteria would exceed that of phage in natural populations. How common that is in natural communities is unclear as in marine settings it is thought that the density of phage far exceeds that of the bacteria (29, 30), though recent studies have questioned how high this ratio actually is (31). There is another caveat in that we have restricted our study to human samples because of the clinical relevance. Other studies by colleagues have examined the phage and *E. coli* interactions in the microbiome of other animals such as horses (32). Despite these caveats, we believe the mechanism proposed here happens not only in the gut but more broadly in nature. We cannot say definitively how pervasive these observations are, but this question of generality could be answered by other investigators with culture-independent approaches.

We expect that the approach developed in this study also has broader utility for translational phage investigation. If phages are to be developed as therapeutics or as components of therapeutic consortia, we expect that this will require complementary culture-based and culture-independent techniques that include experimental isolation, characterization, and strain-resolved analysis in the context of complex communities.

The increasing incidence of antibiotic-resistant infections in recent years has resurrected interest in using phage to treat bacterial infections. When it comes to utilizing bacteriophages therapeutically, one would expect a tradeoff between this generalized resistance mechanism and bacterial virulence. This phenomenon was illustrated in a classical study by Smith and Huggins, where they utilized a phage targeting the K antigen of pathogenic *E. coli* (*33*). Interestingly, the mechanism of resistance explored here, the O-antigen, is known to be the target of some bacteriophages (34). If the results presented here for *E. coli* are supported in other *Enterobacteriaceae*, we would expect difficulties in isolating phage capable of lysing specific bacterial isolates. This anticipated result is echoed by investigators working on isolating phage for therapy where the rates of finding phage for an isolate are less than 50% for some pathogens (35–37). Here we provide one explanation for why this could be the case: that is, resistance associated with production of the O-antigen.

## Materials and Methods

### Fecal Microbiota Transplant Doses

Healthy human participants were recruited and provided informed consent to participate in a stool donor protocol approved by the Emory University Institutional Review Board (Protocol 00112302, PI Woodworth). Donors completed health and behavior questionnaires, blood, urine, and stool laboratory testing to screen for potential pathogens as described in the PREMIX trial (38). Each stool specimen from a single donor was divided by weight with 50-100g of stool in a Seward filter bag and suspended in 250 mL USP sterile saline using a Seward benchtop stomacher at 200 rpm for 30 seconds twice. After removing the filter bag, the remaining suspension of stool in saline was transferred back to the 250mL saline bottle and delivered fresh without freezing for *E. coli* and phage isolation as described below. Additional aliquots with and without 10% glycerol by starting volume were prepared and stored at −80 °C.

### Bacterial Strains

*E. coli* C was obtained from Marie-Agnès Petit at INRAE in Joy-en-Josas, France. MG1655 L9 and L5 were obtained from Douglas Browning at Aston University in Birmingham, UK (39). Briefly, these constructs were made by cloning either the *rfb* cluster or *wwbL* into isogenic MG1655. Wild *E. coli* were isolated from fecal microbiota transplantation doses as described below.

### Bacteriophages

Phages Lambda^VIR^, T2, T3, T4, and T7 were acquired from the Levin Lab’s phage collection. Wild phages were isolated from fecal microbiota transplantation doses as described below.

### Growth Media and Conditions

Unless otherwise noted, all bacteria were cultured in Lysogeny Broth (LB) at 37 °C. Lysogeny broth (LB) prepared according to manufacturer instructions (Difco, 244620). LB soft agar made with 0.7% w/v agarose, and LB plates made with 1.6% agarose, LB phage plates prepared as LB plates supplemented with 20 mM CaCl_2_.

### Isolating Bacteria from the FMTs

Serial dilutions of FMT in 0.85% NaCl solution was followed by plating on Davis Minimal (DM) plates (Sigma-Aldrich, 15758) supplemented with 0.4% w/v lactose (Sigma-Aldrich, L3750). Individual colonies were randomly chosen and picked with a sterile stick and simultaneously streaked onto an EMB plate and a DM plate. Both were incubated at 37 °C overnight. Isolates colored green on the EMB plate were picked from their its respective minimal plate and serially streaked from the DM lactose plate onto another DM lactose plate and incubated at 37 °C overnight. Each isolate was labeled, stored, and frozen.

### Isolating Phage from the FMTs

0.1 mL of FMT suspension was inoculated into 10mL LB broth with ∼1e^7^ CFU/mL *E. coli* C. These flasks were grown with shaking at 37 °C overnight. The next day, samples were centrifuged and filtered through a 0.22-micron filter (Basix, 13-1001-06). These rough lysates were then plated for single plaques. Individual plaques were picked with a sterile stick and resuspended as described above to obtain clonal phage lysates.

### AAI heatmap

We identified the closest relatives to the genomes of this study (query genomes) using whole-genome searches against public genome databases. The final set of genomes used in subsequent analyses consisted of 98 genomes – including the 9 query genomes, 34 genomes from the IMG/VR v4 database (40), and 55 genomes that are publicly available on NCBI. Prodigal v2.6.3 (41) was used for gene prediction with default settings, and pairwise all versus all Average Amino acid Identity (AAI) was calculated using the aai.rb script from the Enveomics collection (42) using default settings except for decreasing the number of minimum hits to 20. The final heatmap was constructed using the pheatmap package in R. Accession numbers of the public genomes are labelled in the figure.

### Core gene tree

The subset of genomes that make up the major cluster in the AAI heatmap were used for gene clustering at 30% identity with MMseqs2 v 13.4511 (43). The resulting clusters were used to identify the core genome – the genes that are found in 55/60 (∼90%) of the genomes. The 32 core genes were extracted using seqtk-1.4 and individual gene alignments were generated using MUSCLE v3.8.31 (44) with default settings. Each alignment was used to generate a maximum likelihood gene tree in MEGA11 (44) with default parameters. These gene trees were concatenated and passed into Astral 5.7.8 (23) to generate the final consensus tree as Astral has a built-in algorithm to deal with any missing data in the input gene trees. The final tree was visualized and annotated in iTol (45) and is shown in Figure 4B. The tree was rooted using *Escherichia virus vB_eco_mar005P1*. The additional two genomes used as outgroups to root this tree were added manually.

### Phage Susceptibility Testing

∼1e^6^ of each phage lysate was spotted onto lawns of the bacteria using the double-layer soft agar technique as described in (46). Phage lysates were diluted 1:100 prior to spotting to minimize lysis from without.

### Sampling Bacterial and Phage Densities

Bacteria and phage densities were estimated by serial dilutions in 0.85% NaCl solution followed by plating. The total density of bacteria was estimated on LB (1.6%) agar plates. To estimate the densities of free phage, chloroform was added to suspensions before serial dilutions and double-layer soft agar enumeration.

### Serial Transfer Experiments

All serial transfer experiments were carried out in 10 mL LB broth cultures grown at 37 °C with vigorous shaking. The cultures were initiated by 1:100 dilution from 10 mL overnight cultures grown from single colonies. Phage was added to these cultures to reach the initial density of∼10^7^ PFU/mL. At the end of each transfer, 0.1 mL of each culture was transferred into flasks with fresh medium (1:100 dilution). Simultaneously, 0.1 mL samples were taken for estimating the densities of CFUs and PFUs as described above.

### Estimation of the Rate of Spontaneous O Antigen Loss

Overnight cultures of *E. coli* K-12 strain MC4100 *wbbL*^+^ (MG873) in which O-antigen synthesis is genetically restored were diluted and plated onto either LB agar (total colony forming units (CFU)) or LB agar supplemented with 175 μg/mL of vancomycin (vancomycin resistant CFU). To determine the proportion of vancomycin-resistant colonies that lost the ability to produce O-antigen, 700 Vancomycin-resistant colonies (from 7 independent experiments) were then screened for sensitivity to Ffm phage that require loss of O-antigen. The rate of O-antigen loss was defined as the number spontaneous vancomycin-resistant CFU divided by the of Ffm-sensitive colonies, adjusted for the rate of Ffm-sensitivity in the vancomycin-resistant colonies. Alternatively, serial dilutions of MG1655L9 were plated on LB plates with 175 μg/mL of vancomycin. 100 surviving colonies from each replicate were screened for sensitivity to the phage P1-sensitivity to which indicates a loss of the O-antigen.

### Phage Suceptibility Due to O Antigen Loss in FMT-derived *E. coli*

Serial dilutions of the FMT-derived *E. coli* were plated on LB plates with containing 3x the ancestoral MIC of vancomycin. Surviving colonies from each replicate were screened for sensitivity to the phage P1-sensitivity to which indicates a loss of the O-antigen-as well as T2, T4, and T7.

### Antibiotics and Their Sources

Streptomycin (S6501) was obtained from Sigma-Aldrich; Tetracycline (T1700) from Research Products International; Gentamicin (BP918-1) from Fisher BioReagents; Chloramphenicol (C5793) from Sigma-Aldrich; Ciprofloxacin (A4556) from AppliChem); Ceftriaxone (C5793) from Sigma-Aldrich; Meropenem (QH-8889) from Combi-Blocks; Azithromycin (3771) from Tocris; Fosfomycin (P5396) from Sigma-Aldrich; and Colistin (C4461) from Sigma-Aldrich. All antibiotic Sensi-Discs were obtained from Becton Dickinson.

### Invasion When Rare Experiment

E. coli MG1655 and E. coli MG1655L9 were co-cultured in flasks at a ratio of 1:1000 either with or without phage T7. Bacterial and phage densities were estimated at 0 and 24 hours.

### O Antigen Antisera Assay

*E. coli* O-antigen-specific antisera was obtained from SSI Diagnostica and used as detailed in their protocol for the slide agglutination assay. Each strain was tested with the O antisera which corresponded to its predicted O-antigen type to confirm expression of this gene cluster. A known O-antigen negative lab strain (MG1655) and its construct bearing the O-antigen (MG1655L9) were used as negative and positive controls respectively.

### Whole-Genome Sequencing

Samples were sent to MIGS (Pittsburgh, USA) as bacterial colonies grown on an agar plate or as sterile phage lysates for extraction and Illumina sequencing. Individual colonies and lysates were extracted per the manufacturer’s protocol using the Zymo DNA miniprep bead beating lysis kit. Sample libraries were prepared for Illumina sequencing using Illumina’s DNA Prep kit and IDT 10bp unique dual indices and sequenced on an Illumina NovaSeq 6000, producing paired end 151bp reads. Demultiplexing, quality control, and adapter trimming was performed with bcl-convert (v4.1.5). Illumina reads were quality filtered using Trimmomatic (47). *E. coli* C host sequences were depleted from phage lysate sequencing data by mapping to a closed *E. coli* C genome with bowtie2 (48). After trimming and host decontamination (for phage lysates), remaining reads were and assembled *de novo* using SPAdes v3.13 (49). Pairwise comparisons of average nucleotide identity on the assembled genomes were performed with the Mashmap method using fastANI v1.32 as in (50). Gene sequences were predicted with Prodigal v2.6.3 (41) and annotated with Prokka v1.14.6 (51). O and H antigens were predicted *in silico* with Ectyper v2.0.1 (52). Multilocus sequence analysis (MLSA) typing was performed using mlst v2.23 (Seemann, https://github.com/tseemann/mlst) with the Achtman seven-locus scheme and the PubMLST database (53). ANI values were converted to pairwise distances (100 − ANI) and used to construct a neighbor-joining phylogenetic tree with the ape package v5.7 in R v4.3 (54). The tree was midpoint-rooted using phytools v2.0 (55), and negative branch lengths were set to a minimum of 0.0001. The tree was visualized as a circular phylogram using ggtree v3.10 and ggtreeExtra v1.12 (56, 57), with concentric annotation rings indicating donor identity, predicted O-antigen serotype, and MLSA sequence type.

### Numerical Solutions (Simulations)

For our numerical analysis of the coupled, ordered differential equations presented in the Supplemental Text, we used Berkeley Madonna with the parameters presented in Supplemental Table 3. Copies of the Berkeley Madonna programs used for these simulations are available at www.eclf.net.

### Statistical Analysis

Statistical tests of difference in proportions of phage plaquing results tabulated as a contingency table were performed with the chisq.test function in stats package version 4.3.0 in R version 4.3.0 using the R studio interface version 2023.06.0. Differences in phage densities across host strains in serial transfer experiments were assessed using a linear mixed-effects model fit by restricted maximum likelihood (REML) with the lme function in the nlme package, with log₁₀-transformed PFU/mL as the response, strain, transfer day, and their interaction as fixed effects, and biological replicate as a random intercept. Post-hoc pairwise comparisons between strains were performed on estimated marginal means with Bonferroni correction for multiple comparisons. Differences in bacterial densities and frequencies between conditions in the invasion when rare experiment (Supplemental Figure 7) were assessed using two-sample t-tests on log₁₀-transformed densities. Resulting p-values <0.05 were considered statistically significant.

## Supporting information

All Supplemental Materials

## Author Contributions

Conceptualization: MHW, BRL, BAB

Methodology: MHW, BRL, BAB, JF, CEB, KM, MG

Software: MHW, KTK, BRL

Validation: MHW, BRL, BAB, KBB, MGH

Formal Analysis: MHW, BRL, BAB, KBB, MGH, JN, KM

Investigation: BAB, TGG, KBB, MGH, JF, DAG, CEB, JN, KM

Resources: MHW, BRL

Data Curation: MHW, BRL

Writing - Original Draft: MHW, BRL, BAB, KBB, DAG, KTK

Writing - Review and Editing: MHW, BRL, BAB, TGG. KBB, MGH JF, DAG, KTK, CEB, KM, MG

Visualization: BAB, TGG KBB, MGH, CEB, MG

Supervision: MHW, BRL, MG

Project administration: MHW, BRL

Funding acquisition: MHW, BRL, MG

## Acknowledgments

We thank Dalia Gulick, Amanda Strudwick, Amalia Cruz, and Candace Miller for their help obtaining and processing the FMTs. We also thank the other members of the Levin Lab for their collaboration and discussion, in particular Andrew Smith, Josh Manuel, Thomas O’Rourke, Ingrid McCall, and Eduardo Roman. We would like to thank BioRender for their software which was used to create the schematic figure. We are particularly grateful to Andrey Letarov for his comments on the BioRxiv version of this manuscript and discussion, which improved this report.

## Data Availability

The Berkeley Madonna program used for the simulations are available at www.eclf.net. All raw sequence data for the *E. coli* and phage genomes shown have been deposited in the sequence read archive (SRA, NCBI, Bethesda, MD, USA) and can be found with bioproject accession PRJNA1028583.

