## Supplementary material for "Enteric Populations of *Escherichia coli* are Likely to be Resistant to Phages Due to O-Antigen Expression": All Supplemental Materials

**This PDF file includes:**

Supplemental Text

Figures S1 to S7

Tables S1 and S3

**Supplemental Text**

In the following, we use a mathematical-computer simulation model to illustrate how populations dominated by phage-resistant, O antigen-bearing bacteria will maintain populations of lytic and temperate phages.

*Model of the population dynamics of resistant O antigen bearing bacteria and lytic phage*

There are two populations of bacteria, one bears an O antigen, O, that masks the phage receptor, and the other is a variant of the O antigen-bearing bacteria, S, for which the receptor is available for the phage to adsorb. There is a single population of lytic phage, P. The variables O, S, and P are the designations of these populations and their densities, cells and particles per ml for the bacteria and phage, respectively.

The bacteria replicate at a rate proportional to their maximum growth rates, respectively v_O_ and v_S_, and a hyperbolic function of the concentration of a limiting resource, r µg/ml.
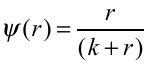
 where k is the resource concentration when the growth rate is half its maximum value, the Monod constant (1). For our analysis of the properties of this model, we use the same value of k for both O and S. The limiting resource is taken up at a rate proportional to the sum of the product of the densities of the populations, their maximum growth rates, the Monod function, ψ(r) and a conversion efficiency parameter, e µg per cell. The phage infects the bacteria at a rate equal to the product of the densities of bacteria and phage and rate parameter, δ_O_ and δ_S_ per ml, ml attacks per phage particle per cell per hour with, δ_O_ << δ_S_. Each infected bacterium produces β phage particles. The O population transitions to S, and the S transition to O at a rate of µ_OS_ and µ_SO_ per cell per hour, respectively. To account for the decline in the physiological state of the bacteria as the resource decline, we assume the rates of phage infection and transition between states are proportional to ψ(r) (2, 3).

With these definitions and assumptions, the rates of change in the concentration of the limiting resource and densities of the bacteria and phage are given by,

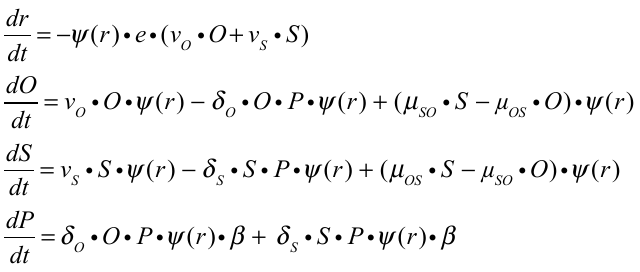

To solve these equations and those that follow for the temperate phage, we use Berkeley Madonna. In these numerical solutions, simulations, we assume the populations are maintained in serial transfer culture. Every 24 hours, the populations of bacteria and diluted by a factor of 1/100, and 1000µg/ml of fresh resource is added.

In Figure 4, we present the results of simulations with three rates of transition between O and S, µ_OS_ and µ_SO_ per cell per hour.

If the rate of transition between the sensitive, S, and resistant O antigen-bearing state is great enough, the phage is maintained in a population dominated by resistant O antigen-bearing bacteria. In the above simulations, we are assuming sensitive and resistant, S and O, bacteria have the same growth rate. The conditions for the maintenance of the phage would be greater if the O antigen mediated resistance had a fitness cost, v­_O_ < v_S_.

*Model of the population dynamics of resistant O antigen bearing bacteria and temperate phage*

There are two populations of lysogenic bacteria, one bearing the O antigen and one without the O antigen, with designations and densities OL and L, respectively. There is a single population of free temperate phage of density PT particles per ml. The phage can adsorb to L, with a rate constant δ, but cannot adsorb to the OL. Temperate phages that adsorb to lysogens are lost. With rates of ind_OL_ and ind_L_, the OL and L lysogens are induced and produce β_OL_ and β_L_ phage particles, respectively, with the induced OL and L dying. As with the lytic phage model, we assume that all rates are proportional to the concentration of the limiting resource by a Monod function, $\psi\left( r \right)=\frac{r}{(r+k)}$, and the limiting resource is taken up at a rate proportional $\psi\left( r \right)$, their growth rates and densities, and a conversion efficiency parameter, e µg/per cell. We neglect the loss of the prophage by the OL and L lysogens.

With these definitions and assumptions, the rates of change in the densities of bacteria are given by,

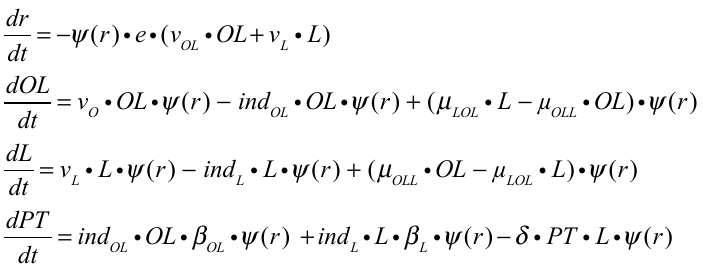

In Supplemental Figure 3, we present the results of simulations of the population dynamics of temperate phage and lysogens that mask the receptor and do not adsorb the phage, OL, and lysogens with the receptor open to adsorption by PT, L.

In all cases bacterial populations are dominated by “resistant”, O antigen bearing lysogens, substantial densities of free temperate phage are maintained.

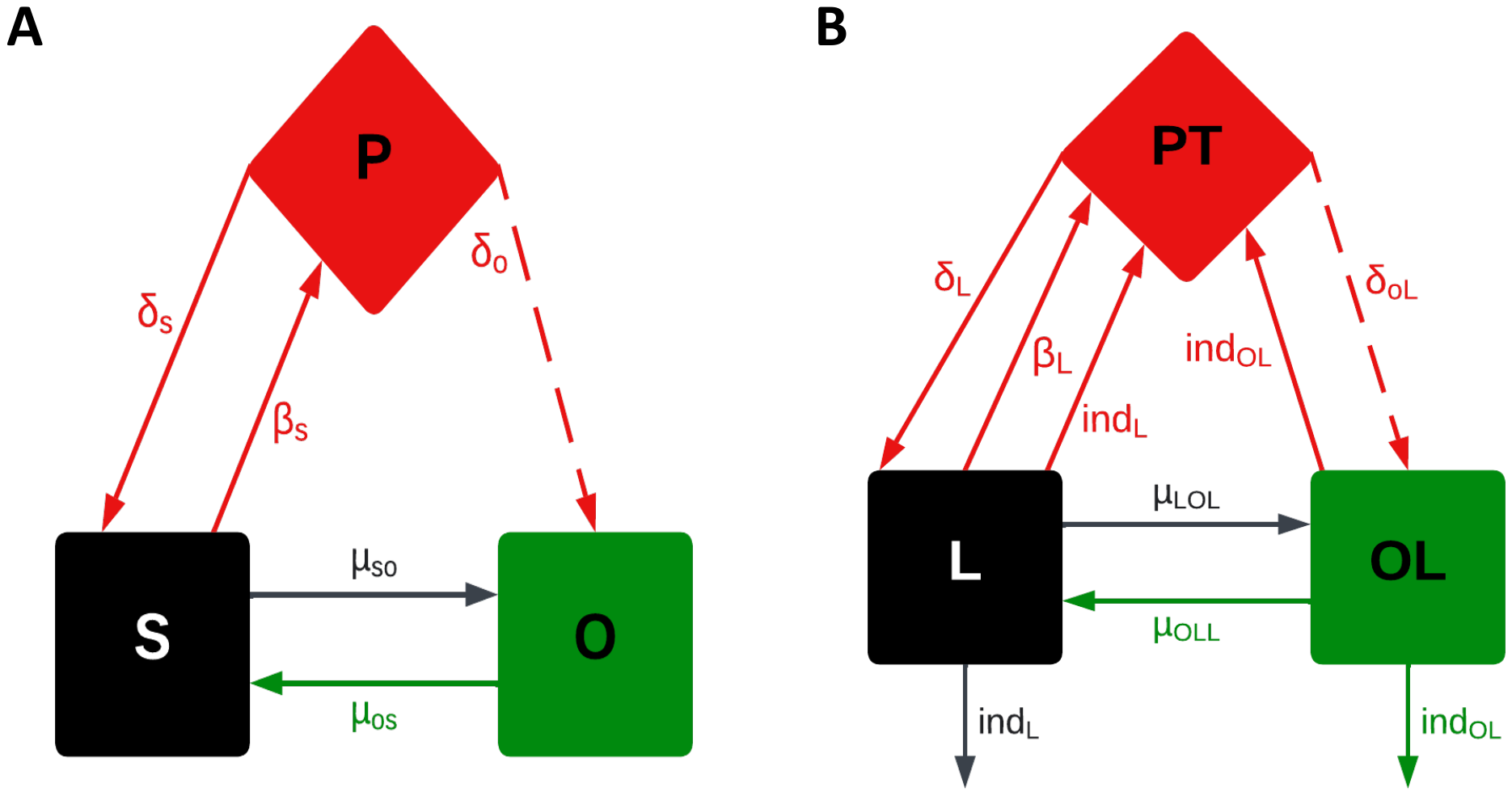

**Figure S1**. **Models of the population and evolutionary dynamics of O antigen mediated phage resistance.** See the supplemental text for a description of the model and Supplemental Table 1 for the definitions and dimensions of the parameters and the values used in our numerical solutions to the equations and simulations. **A)** There is a population of lytic phage, P; a population of phage-sensitive bacteria, S; and a population of phage-resistant, O antigen producing cells, O. **B)** There is a population of temperate phage, PT; a population of immune lysogens, L; and a population of phage-resistant, O antigen producing lysogens, OL.

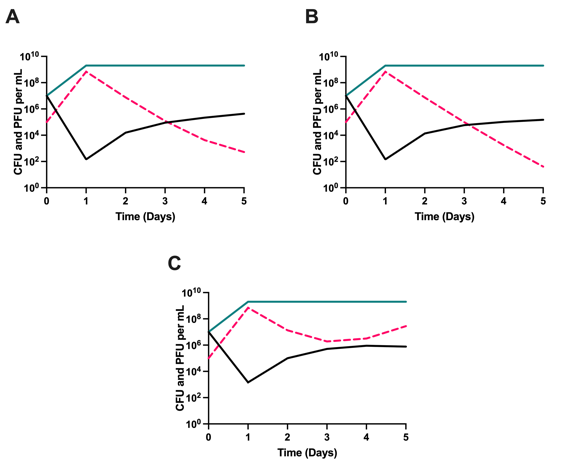

**Figure S2. The effect of fitness cost and unequal transition rates on phage maintenance.** Simulation results, changes in the initial and end of transfer densities of bacteria and phage in serial transfer culture. 1000µg/ml of the limiting resource is added at each transfer. Standard parameters unless otherwise stated, v_O_ = v_S_ = 2.0 per cell per hour, e= 5x10^-7^ µg/cell, k=1.0, β=50, δ_O_=0, δ_S_ – 2x10^-7^ per phage per cell per hour, µ_OS_=µ_SO_=10^-5^ per cell per hour. **A-** v_O_= 1.8 per cell per hour, v_S_ = 2.0 per cell per hour. **B-** µ_OS_= 10^-5^ per cell per hour, µ_SO_=10^-4^ per cell per hour. **C-** µ_OS_= 10^-4^ per cell per hour, µ_SO_=10^-5^ per cell per hour. O antigen expressing cells- turquoise, Cells not expressing the O antigen-black, and Phage – pink.

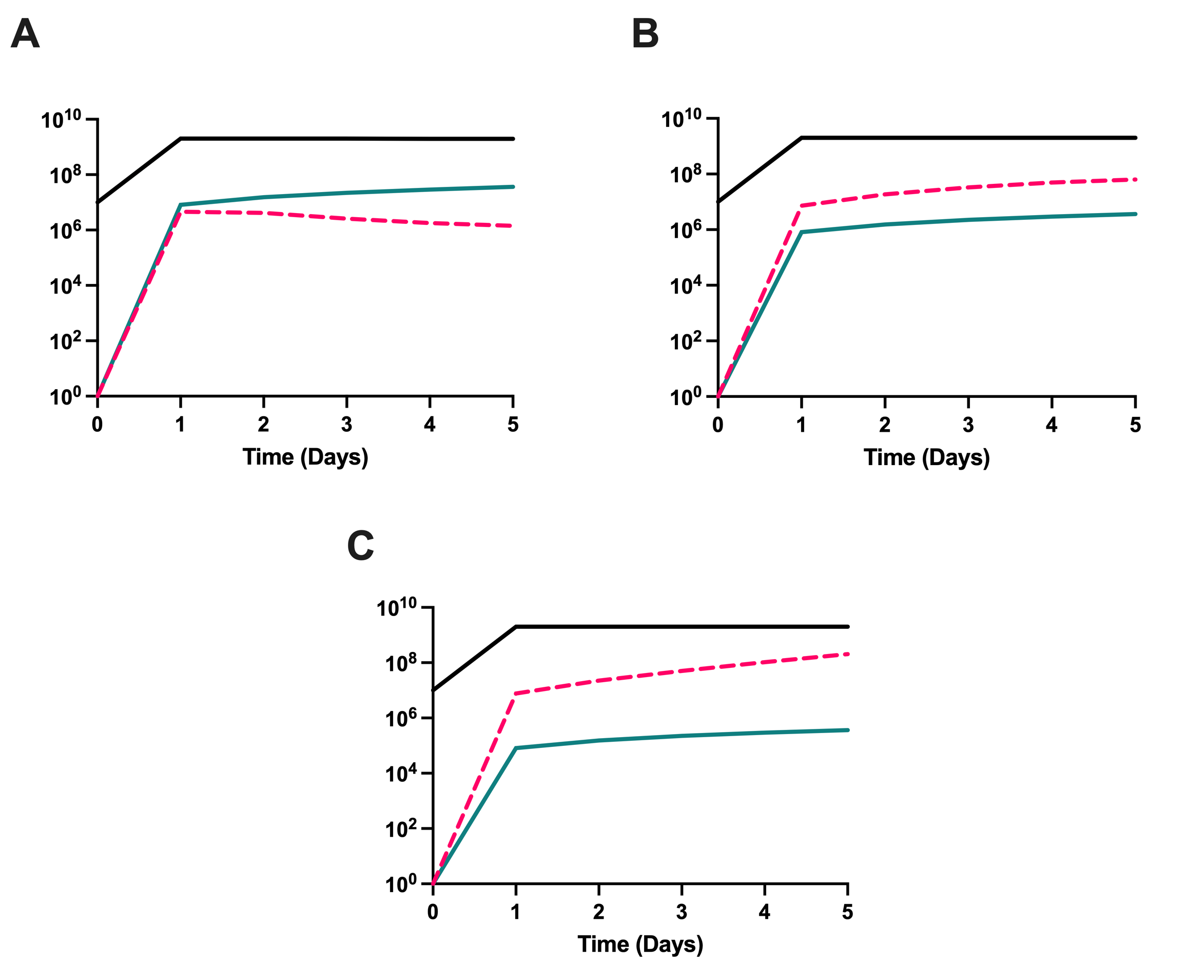

**Figure S3. Predicted serial transfer dynamics of O antigen mediated resistance and a temperate phage.** Simulation results, changes in the initial and end of transfer densities of bacteria and phage in serial transfer culture. 1000µg/ml of the limiting resource is added at each transfer. Standard parameters, v_O_ = v_S_ = 2.0 per cell per hour, e= 5x10^-7^ µg/cell, k=1.0, β=50, δ_O_=0, δ_S_ – 2x10^-7^ per phage per cell per hour. **A-** µ_OS_=µ_SO_=10^-5^ per cell per hour. **B-** µ_OS_=µ_SO_=10^-4^ per cell per hour, and **C-** µ_OS_=µ_SO_=10^-3^ per hour. O antigen expressing lysogenic cells- turquoise, Lysogenic cells-black, Phage – pink.

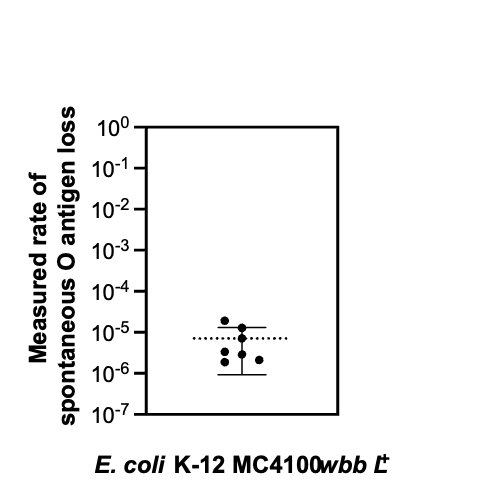

**Figure Sn. Estimated rates of spontaneous O antigen loss in E. coli cultures.** Cultures of MC4100 wbbL+ O antigen producing strain were grown in LB to saturation. CFU in the culture were quantified as total CFU. Vancomycin-resistant subpopulations were selected on LB supplemented with vancomycin (150 mcg/ml) and enumerated to estimate their rate in the total population. In each of seven independent experiments, 100 vancomycin-resistant colonies were screened for sensitivity to Ffm phage, confirming the loss of O antigen as the cause of vancomycin resistance. The rate of Ffm sensitivity was used to correct the vancomycin-resistance rate in the total population to give the presented estimated rate of cells that have undergone spontaneous O antigen loss within a population. Mean (dotted line) with 95% CI range is presented.

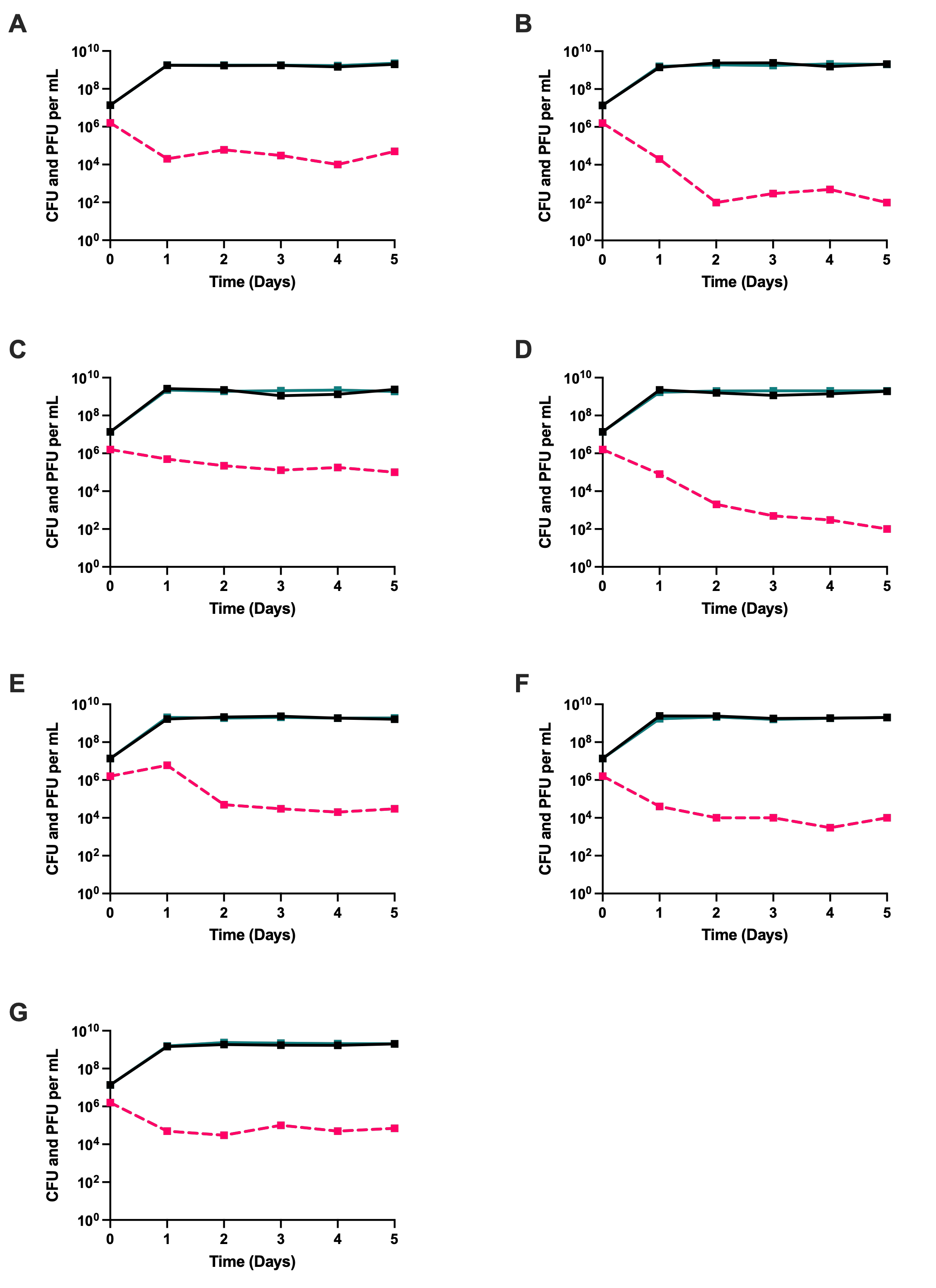

**Figure S5. Serial transfer replicates of one of each serotypes of the FMT-isolated *E. coli*.** Seven *E. coli* isolates serially transferred for five days in LB broth with the phage T7. Shown in solid lines are bacterial densities with the black line representing a phage-free control. The broken lines present phage densities. **(A)** A16. **(B)** B13. **(C)** B14. **(D)** B22. **(E)** C11. **(F)** C15. **(G)** D6.

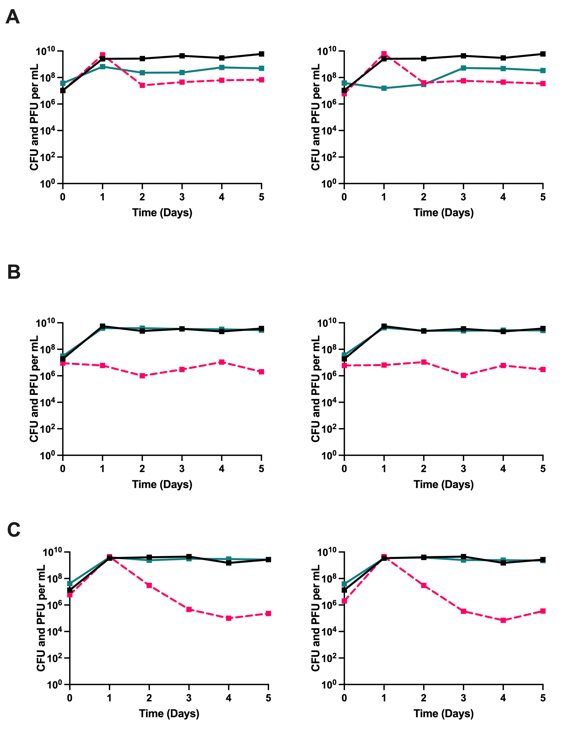

**Figure S6. Serial transfer replicates of T7 with MG1655 lacking and expressing the O antigen.** Three variants of *E. coli* MG1655 serially transferred for five days in LB broth with the phage T7. Shown in solid lines are bacterial densities with the black line representing a phage free control. The broken lines present phage densities. Shown side by side two of three biological replicates. **(A)** Wild type MG1655 not expressing the O antigen. **(B)** MG1655L5 expressing a medium amount of the O antigen. **(C)** MG1655L9 expressing a high amount of the O antigen

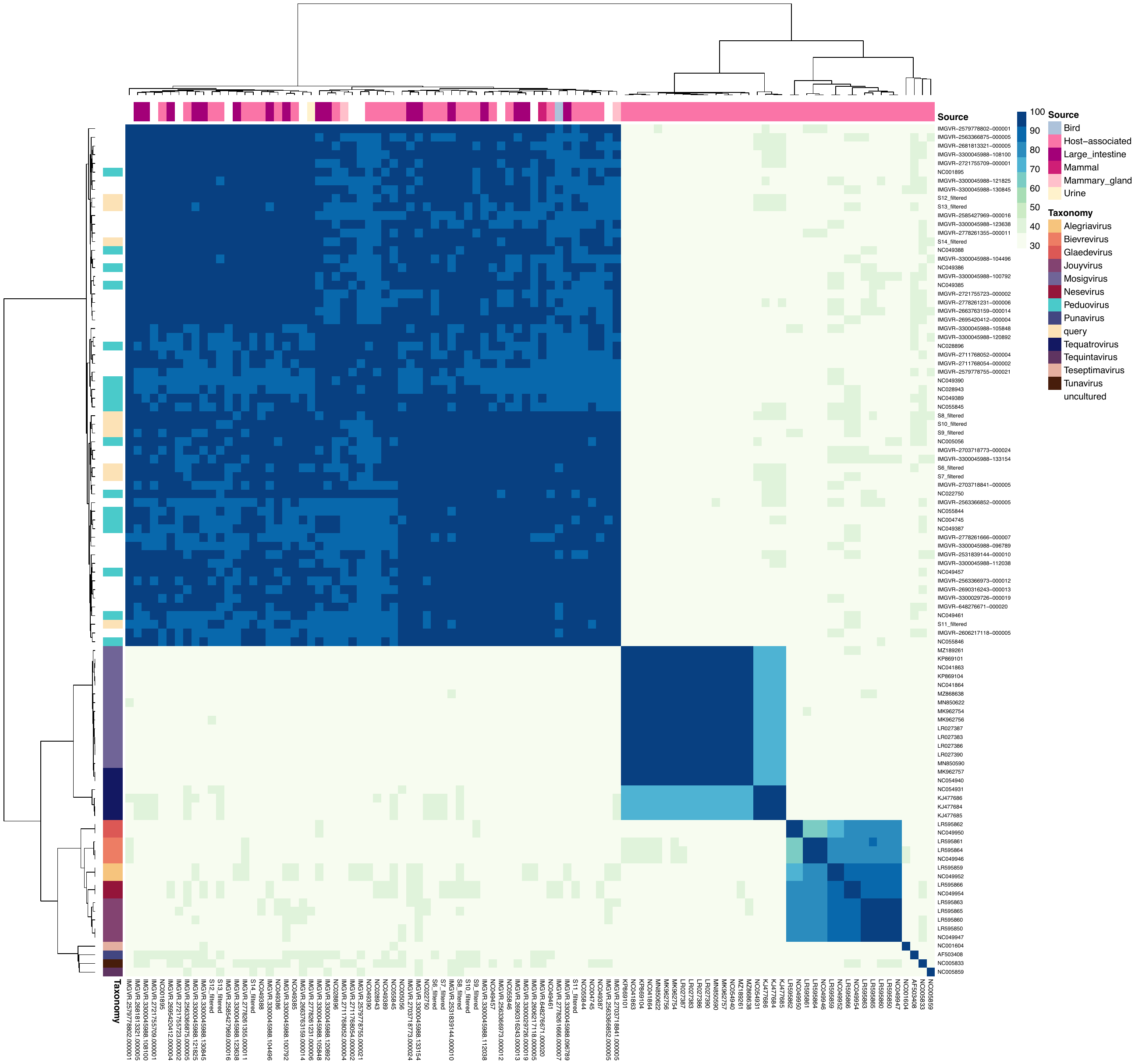

**Figure S7. Heatmap of average amino acid identities (AAIs) among genomes of this study and selected reference genomes from public databases.** AAI was estimated based on BLAST+ whole-genome comparisons using a minimum alignment identity of 20% and at least 20 genes shared between two genomes. Genomes of this study were clearly placed in the major AAI cluster that included all *Peduovirus* genomes from IMG/VR and several close relatives from NCBI.

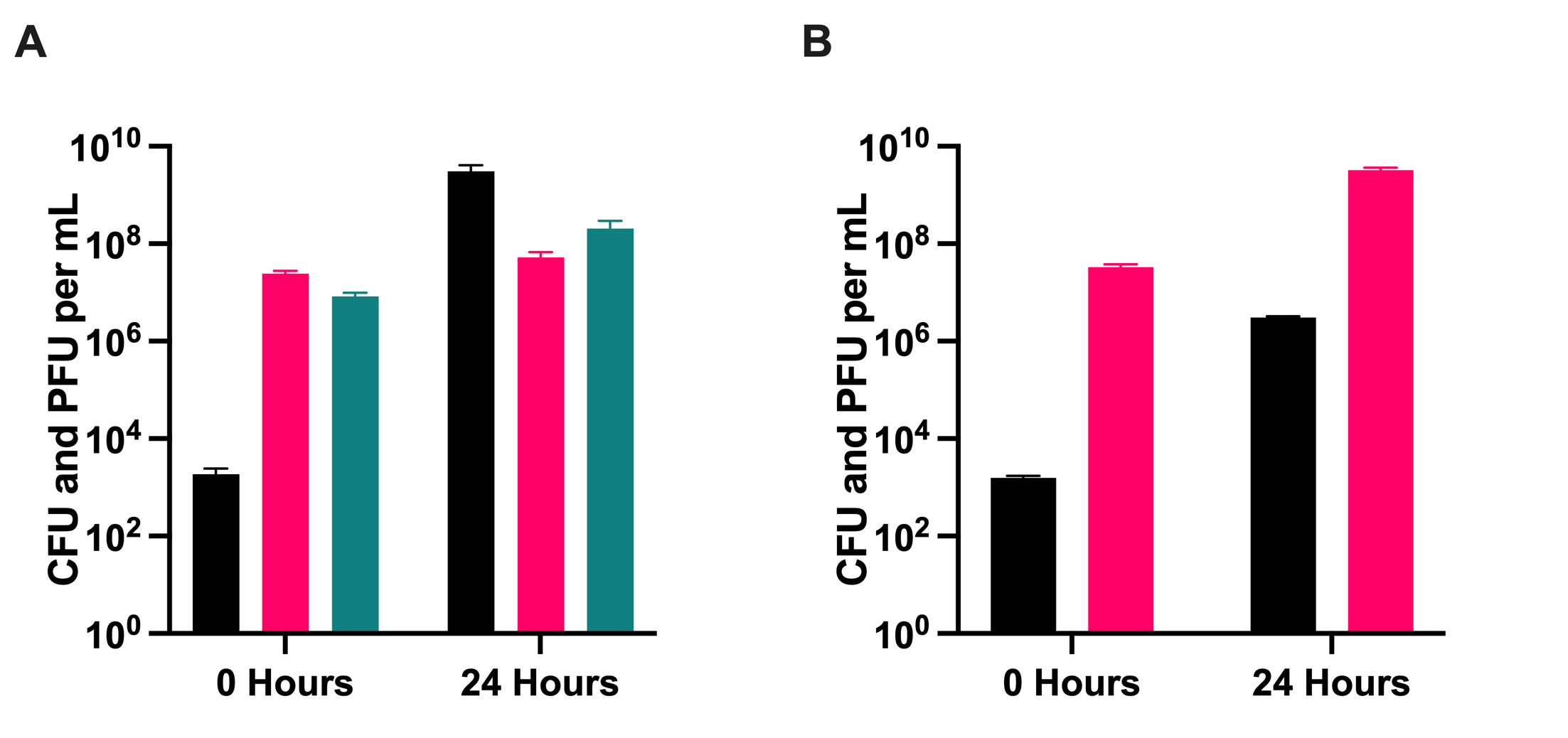

**Figure S8. Invasion when rare experiment of O antigen expressing cells into a population of O antigen deficient cells.** Shown are the results of an experiment initiated with **(A)** 10^3^ O-antigen expressing cells, 10^7^ O-antigen deficient cells, and 10^7^ lytic phage or **(B)** 10^3^ O-antigen expressing cells, 10^7^ O-antigen deficient cells, and no phage. Shown are mean and standard deviations of three biological replicates. O antigen expressing cells are in black, O antigen deficient cells in fuchsia, and a lytic phage in turquoise. In the presence of phage T7, O-antigen expressing cells increased from ~0.008% to 98.1% of the total population within 24 hours, compared to 0.09% without phage (t-test p = 2.9 × 10⁻⁵), indicating phage-mediated selection for O antigen expression.

**Table S1:** Summary of lysate spotting experimental results.

|  | **None (N=392)** | **Turbid (N=85)** | **P-value** |
| --- | --- | --- | --- |
| **FMT_cross** |  |  |  |
| Allogeneic | 304 (77.6%) | 56 (65.9%) | 0.0334 |
| Autologous | 88 (22.4%) | 29 (34.1%) |  |
| **O1** |  |  |  |
| 17 | 143 (36.5%) | 46 (54.1%) | <0.001 |
| 74 | 27 (6.9%) | 27 (31.8%) |  |
| 132 | 9 (2.3%) | 0 (0%) |  |
| 1 | 26 (6.6%) | 1 (1.2%) |  |
| 8 | 9 (2.3%) | 0 (0%) |  |
| 16 | 9 (2.3%) | 0 (0%) |  |
| 167 | 9 (2.3%) | 0 (0%) |  |
| None predicted | 160 (40.8%) | 11 (12.9%) |  |
| **O2** |  |  |  |
| 1 | 9 (2.3%) | 0 (0%) | <0.001 |
| 16 | 9 (2.3%) | 0 (0%) |  |
| 44 | 76 (19.4%) | 14 (16.5%) |  |
| 74 | 14 (3.6%) | 13 (15.3%) |  |
| 77 | 67 (17.1%) | 32 (37.6%) |  |
| 132 | 9 (2.3%) | 0 (0%) |  |
| None predicted | 208 (53.1%) | 26 (30.6%) |  |
| **O3** |  |  |  |
| 17 | 64 (16.3%) | 17 (20.0%) | 0.754 |
| 44 | 13 (3.3%) | 5 (5.9%) |  |
| None predicted | 315 (80.4%) | 63 (74.1%) |  |
| **O4** |  |  |  |
| 44 | 10 (2.6%) | 8 (9.4%) | 0.0146 |
| 77 | 54 (13.8%) | 9 (10.6%) |  |
| None predicted | 328 (83.7%) | 68 (80.0%) |  |
| **Distinct O Antigen** |  |  |  |
| Mean (SD) | 1.15 (1.16) | 1.67 (1.00) | <0.001 |
| Median [Min, Max] | 1.00 [0, 3.00] | 2.00 [0, 3.00] |  |
| **H** |  |  |  |
| 4 | 9 (2.3%) | 0 (0%) | <0.001 |
| 5 | 9 (2.3%) | 0 (0%) |  |
| 7 | 41 (10.5%) | 4 (4.7%) |  |
| 8 | 9 (2.3%) | 0 (0%) |  |
| 9 | 9 (2.3%) | 0 (0%) |  |
| 18 | 162 (41.3%) | 54 (63.5%) |  |
| 19 | 9 (2.3%) | 0 (0%) |  |
| 39 | 27 (6.9%) | 27 (31.8%) |  |
| 45 | 117 (29.8%) | 0 (0%) |  |

Table S2. Phage Sensitivity Induced by O Antigen Loss

| Strain | Serotype | P1 Susceptibility | | T2 Susceptibility | | T4 Susceptibility | | T7 Susceptibility | | Vancomycin  Susceptibility | |
| --- | --- | --- | --- | --- | --- | --- | --- | --- | --- | --- | --- |
|  |  | Initial | Final | Initial | Final | Initial | Final | Initial | Final | Initial | Final |
| A16 | 132 | - | + | - | + | - | + | - | + | 16 | 512 |
| B14 | 1 | - | + | - | + | - | + | - | + | 16 | 256 |
| B13 | 8 | - | + | - | + | - | + | - | + | 16 | 256 |
| B22 | 167 | - | + | + | + | - | + | - | + | 16 | 256 |
| C11 | 17/77/44 | - | + | + | + | + | + | + | + | 16 | 256 |
| C15 | 16 | - | + | - | + | - | + | - | + | 16 | 512 |
| D6 | 74 | - | + | + | + | - | + | - | + | 16 | 128 |

Table S3. Parameter definitions, units, and values used in generating the mathematical models and computer simulations.

| Parameter | Value (dimensions) | Description | Source |
| --- | --- | --- | --- |
| v_O,_ v_S_ | 2.0 (h^-1^) | Maximum growth rates | This paper |
| μ_OS,_ µ_SO_ | 1e^-^5, 1e^-5^, or 1e^-4^ (h^-1^) | Transitions O 🡪 S, S 🡪 O | Chaudhry 2018 (4) |
| δ_O,_ δ_S_ | 2e^-7^ (h^-1^⋅mL^-1^) | Adsorption rate constants | Chaudhry 2018 (4) |
| β | 60 (PFU⋅CFU^-1^) | Burst size | Berryhill 2023 (5) |
| λ | 1e^-2^ | Probability of lysogeny | Berryhill 2023 (5) |
| ind | 1e^-4^ (h^-1^) | Induction rate | Berryhill 2023 (5) |
| k | 1 (µg) | Monod constant | Stewart and Levin 1973 (3) |
| e | 5e^-7^ (µg⋅CFU^-1^) | Conversion efficiency | Stewart and Levin 1973 (3) |
